# Identifying cell-state associated alternative splicing events and their co-regulation

**DOI:** 10.1101/2021.07.23.453605

**Authors:** Carlos F. Buen Abad Najar, Prakruthi Burra, Nir Yosef, Liana F. Lareau

## Abstract

Alternative splicing shapes the transcriptome and contributes to each cell’s unique identity, but single-cell RNA sequencing has struggled to capture the impact of alternative splicing. We previously showed that low recovery of mRNAs from single cells led to erroneous conclusions about the cell-to-cell variability of alternative splicing (1). Here, we present a method, Psix, to confidently identify splicing that changes across a landscape of single cells, using a probabilistic model that is robust against the data limitations of scRNA-seq. Its autocorrelation-inspired approach finds patterns of alternative splicing that correspond to patterns of cell identity, such as cell type or developmental stage, without the need for explicit cell clustering, labeling, or trajectory inference. Applying Psix to data that follow the trajectory of mouse brain development, we identify exons whose alternative splicing patterns cluster into modules of co-regulation. We show that the exons in these modules are enriched for binding by distinct neuronal splicing factors, and that their changes in splicing correspond to changes in expression of these splicing factors. Thus, Psix reveals cell-type-dependent splicing patterns and the wiring of the splicing regulatory networks that control them. Our new method will enable scRNA-seq analysis to go beyond transcription to understand the roles of post-transcriptional regulation in determining cell identity.

---

Transcriptome profiling at a single-cell level has revolutionized our understanding of the continuous biological variation in gene expression that determines a cell’s unique identity (2, 3). Alternative mRNA splicing is a major source of transcriptome variability that plays an important role in determining the identity of a cell (4, 5), but single-cell analyses have not generally distinguished between different transcript isoforms of a gene. This misses a major source of biological variation; the fine-tuned regulation of splicing contributes to many continuous biological processes such as neurogenesis (6, 7), while its misregulation is associated with complex diseases (8–11). Thus, a more complete understanding of gene expression variability between cells and its phenotypic consequence requires an evaluation of changes in splicing and inference of how these changes are regulated.

Despite the enormous progress in computational modeling of cell identity from single-cell gene expression studies (12–16), formidable challenges remain in capturing the impact of alternative splicing (17). A major limitation to this end is the sensitivity with which alternative splicing events can be read from a single cell. Generally, alternative isoforms are distinguished by only a few specific regions of the transcript, which influences accuracy even in bulk level studies. This limitation becomes more acute in single-cell RNA-sequencing data due to low capture efficiency and extensive PCR amplification, which add technical variation and bias. As a result, the observed rates of an exon’s inclusion (described for each cell as the percent of transcripts from its gene in which the exon is present, or Ψ) are greatly distorted, with an inflation of spurious extreme values (1, 17), especially in exons from moderately or lowly expressed genes.

The high observed variance in an exon’s inclusion rate between cells (18, 19), whether due to biological stochasticity or to the technical artifacts resulting from low mRNA capture efficiency (1), can obscure regulated, biologically important splicing changes between different cell types or states. Computational methods have endeavored to reveal these examples amidst the noise. One method used spike-in transcripts to measure and model the variance expected from technical noise and looked for splicing with variance beyond this (20). Several studies have mitigated the impact of technical variance by looking for differences in the mean inclusion rate of an exon between defined groups of cells, expecting that changes in exon inclusion would be noticeable between cell groups as a whole even when technical noise is high (1, 20, 21). The problems of low coverage can be further alleviated by incorporating extra information such as gene sequence and cell type as priors in estimating rates of exon inclusion (22, 23). However, none of these approaches explicitly model the distortion of splicing observations caused by low capture efficiency. Further, methods that require cells to be clustered by condition or sample cannot take full advantage of the insight that single-cell data provides into continuous biological processes. A method that analyzes splicing across a continuous cell population, while considering the impact of low mRNA capture efficiency, has yet to be developed.

Our goal in this work is to confidently identify alternative splicing events that vary between cells, while accounting for limitations in sensitivity and not enforcing any *a priori* stratification of cells into subpopulations or trajectories. Our approach relies on the notion of increasing sensitivity by using information beyond the observed splicing of the specific exon of interest in the cell of interest. To this end, we frame our objective as the detection of alternative splicing events that reflect changes in cell state, as defined by the entire transcriptome. For instance, an alternative exon of interest might be spliced into mRNAs at low levels in stem cells and at increasingly high levels in differentiating cells (Figure 1a). While such an observation may only be supported by a small number of captured mRNA molecules, we posit that its consistency with cell state—as reflected in the similarity of the exon’s observed splicing between similar cells—makes it more likely to represent the underlying biology. As we demonstrate next, this definition does not require dividing the cells into groups by clustering or by labeling cell types, nor does it require an explicit trajectory of cell progression.

**Fig. 1.**
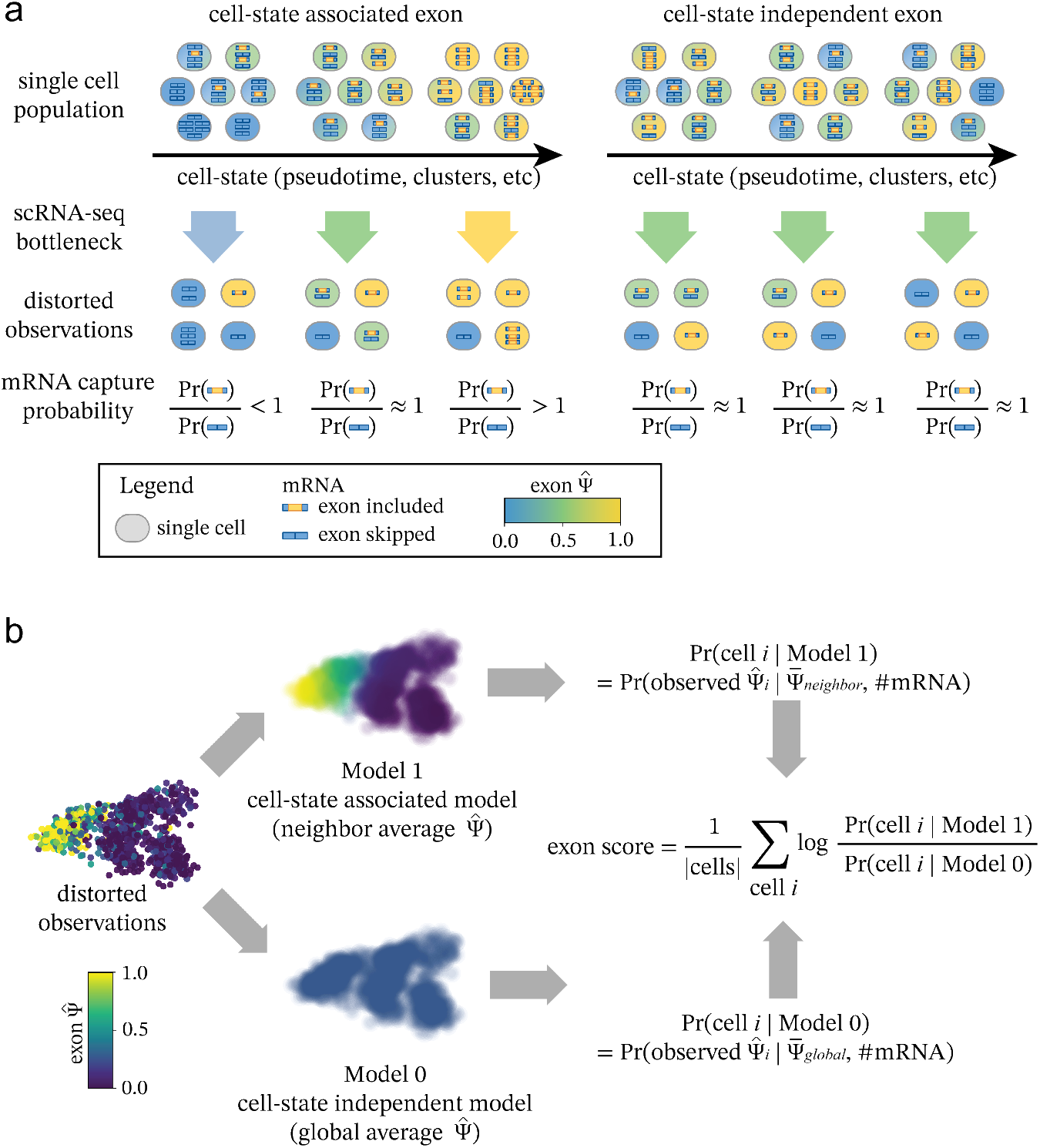
a) Cell-state associated exons change across the phenotypic landscape of a single-cell population. Cell-state independent exons do not change across the phenotypic landscape. Low capture efficiency in scRNA-seq experiments adds additional technical variance depending on the number of captured mRNA molecules. The probability of capturing each alternative isoform depends on the underlying distribution of exons in the single-cell population. b) Psix compares the likelihood of each single-cell observation given two models: Model 1 in which the exon is cell-state associated (probability of the cell’s 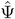 given the average 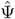 of its *k* nearest neighbors), vs Model 0 in which the exon is cell-state independent (probability of the cell’s 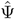 given the average Ψ of all cells in the dataset). Model 1 is more likely for a cell-state associated exon, while the likelihood of both models is similar for a cell-state independent exon.

We formulate these ideas as Psix, a probabilistic method inspired by autocorrelation models that identifies splicing events that are associated with cell state. Psix estimates the likelihood of an exon’s observation in each cell, 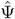, given two models: in the foreground model, we assume that the exon inclusion should be similar to that observed in other, transcriptionally similar cells. In the background model, we assume that the observed exon inclusion reflects sampling noise around a global average (defined separately for each exon) rather than the state of any particular cell. If the exon is associated with cell state, then the first model would be more likely than the second (Figure 1b). We formalize this as a score for each exon that consists of the likelihood ratio between the two models, and assign it an empirical *p*-value through randomizing the location of each cell in the low-dimensional manifold. The evaluation of the two likelihood models requires two things: knowledge of which cells are similar (in transcriptome space), and a model of the distribution of observed exon inclusion values, 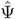, given an underlying unknown Ψ. For the former, we take the similarity to be the distance between cells in a low-dimensional projection based on gene expression profiles (in this study we use a PCA projection of the SCONE normalized gene expression (24); other existing methods can also be used (12–14)). For the latter, we adopt a binomial model of sampling without replacement, reflecting the observation that if few mRNA molecules are captured, the observed exon inclusion rates 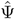 will deviate greatly from the underlying Ψ (1).

We validated our approach using simulated single-cell splicing data from a model of a continuous trajectory of cell states. The simulation allows us to test Psix’s ability to distinguish between two classes of simulated cassette exons: those whose inclusion rates depend on the cell’s position in the trajectory, and those simulated with no change in splicing across the trajectory (i.e., exons reflecting noise properties of this data (1)). First, we simulated gene expression mRNA counts across a continuous trajectory using SymSim (25). For each gene, we then fit a continuous function corresponding to the average underlying Ψ of its cassette exon at a given point of the trajectory: an impulse function for exons with splicing change, and a flat line for exons without splicing change. These classes were assigned randomly to each exon, independently of the simulated mRNA counts of each cell. This underlying Ψ was then used to subsample the mRNA counts to simulate the exon inclusion in a subset of mRNA molecules. Finally, we simulated mRNA capture and short-read, full-coverage sequencing, then subsampled reads to reflect the limited number that cover the splice junctions. This resulted in simulated data that shows similar increases in extreme values as observed in real data (1) (Figure 2a). We tested Psix’s performance on correctly classifying exons as simulated with splicing change (cell-state associated), or as simulated without splicing change (cell-state independent).

**Fig. 2.**
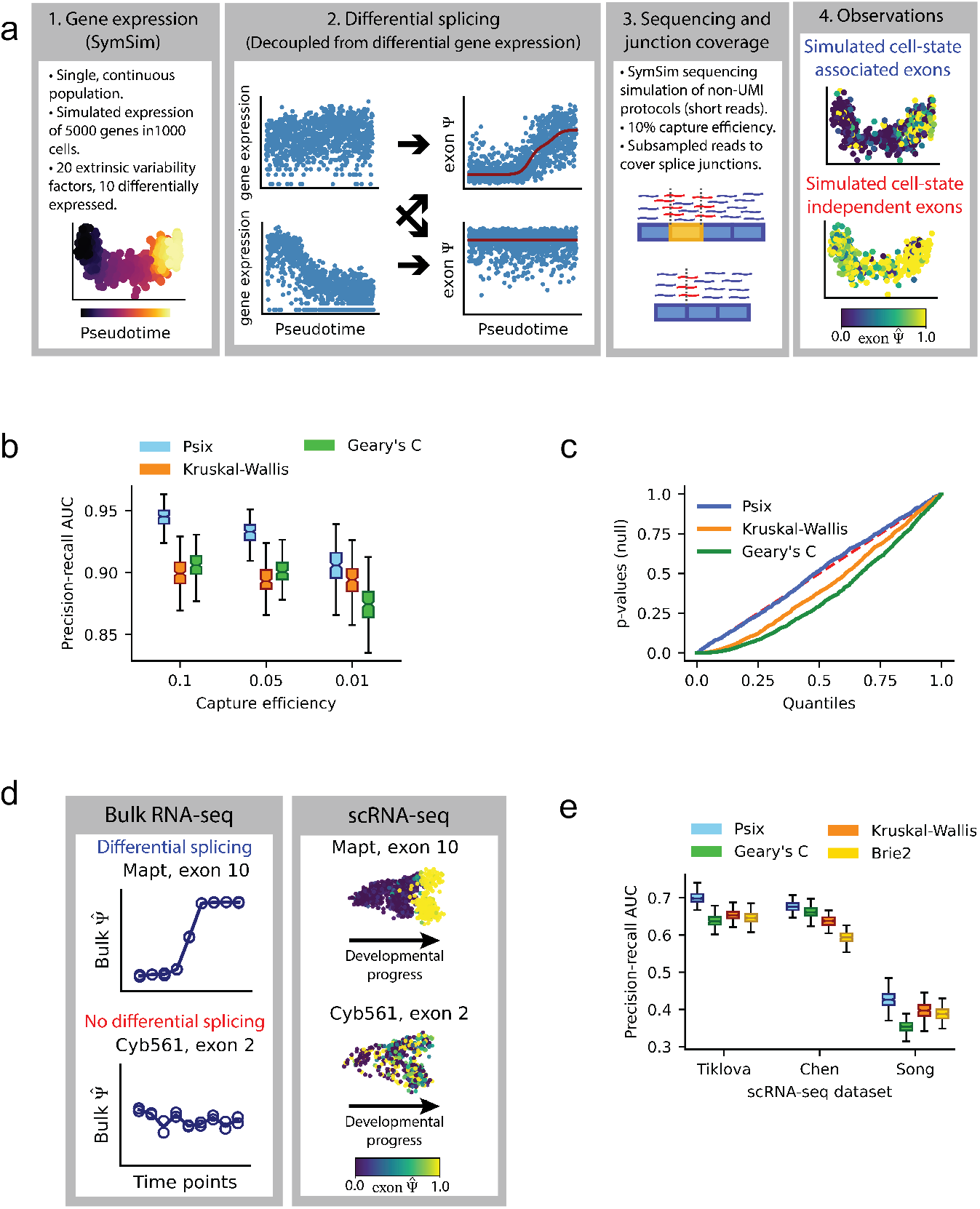
a) Pipeline for simulation of single-cell splicing. Variance in the observations can come from a change in splicing across the cell state (positives), or other sources; e.g., technical noise, or true variance that is not associated with the primary axes of variation in cell state (here, determined by the simulated trajectory; negatives). b) Area under the precision-recall curve showing success of Psix and other methods at identifying exons simulated to have a | ΔΨ | ≥ 0.2, under three different capture efficiencies. c) *p*-value distributions of the negative exons when tested with Psix, Kruskal-Wallis, and Geary’s C. The *p*-values of Psix do not deviate significantly from the uniform distribution. d) Validation strategy for cell-state associated splicing in single cells based on comparable bulk RNA-seq data. e) Area under the precision-recall curve representing the overlap of the exons selected from real scRNA-seq data with the exons from corresponding bulk datasets.

A central feature of Psix is its usage of knowledge of which cells are similar to each other in a continuous phenotype. To test the advantage of this cluster-free approach, we compared Psix’s performance on classifying the simulated exons against a Kruskal-Wallis test that detects differences in the median Ψ between cell clusters (1, 21) (comparing different parts of the simulated trajectory). Another defining characteristic of Psix is its explicit model of the distribution of observed 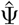 values given an underlying unknown Ψ. To test the contribution of this aspect of our model, we also compared Psix’s results against estimating a Geary’s C, an auto-correlation (cluster-free) test statistic that does not model the number of mRNAs. We found that Psix outperformed these alternative approaches in detecting splicing changes on data simulated at a range of different capture efficiencies (Figure 2b). Psix also outperformed these approaches when simulating a more complex single-cell population with multiple alternative trajectories (Supplementary Figure 1). Importantly, modeling the technical distortion due to low mRNA recovery enabled Psix to avoid excess false positives, unlike other methods (Figure 2c). Furthermore, we note that the relevant splicing events were simulated in a manner that is decoupled from the expression of the respective gene. This allowed us to test Psix’s ability to discriminate between exons simulated with cell-state associated splicing and exons simulated with cell-state independent splicing, irrespective of changes in gene expression.

Next, we sought to test Psix’s ability to identify alternatively spliced exons in real single-cell RNA-seq data. We applied Psix to three different published neurogenesis scRNA-seq datasets. We tested Psix’s ability to identify exons that were observed as differentially spliced in bulk RNA-seq time series datasets that closely matched the scRNA-seq datasets. For instance, a single-cell dataset of embryonic mouse neurons was matched with a bulk RNA-seq time series dataset taken from similar time points, and exons discovered as differentially spliced between time points in the bulk dataset were considered as positives (Figure 2d). We used rMATS (27) to compare each time point in the bulk time series to the first time point, and classified the exons with a significant change (*q*-value ≤ 0.05 and 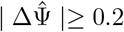) as differentially spliced. We compared Psix against the Kruskal-Wallis and Geary’s C tests as described above. We also compared Psix against an existing approach, BRIE2 (23), which identifies splicing differences between predefined groups of single cells. Psix outperformed all methods in classifying the exons from all three datasets (Figure 2e). (We did not test BRIE2 on our simulations as it requires the actual sequencing data as input, while our SymSim-based simulator generates count matrices directly.)

Because scRNA-seq reports on the biological state of hundreds or thousands of single cells, it has great potential for uncovering regulatory networks that control changes in individual gene expression. Single-cell data have been used successfully to infer transcription regulatory networks (28), but inference of splicing regulation has been very limited, and generally involves pooling predefined clusters of cells (23, 29). Alternative splicing is regulated by a network of splicing factors, and we posited that scRNA-seq data could reveal coordinated changes in co-regulated exons as well as the corresponding changes in expression of the splicing factors that regulate these exons.

To begin reconstructing splicing regulatory networks, we first applied Psix to a dataset of midbrain dopamine neurons across different stages of mouse development (26) (Figure 3a). Psix identified 798 cell-state associated exons over the brain development landscape, including many exons that have been extensively reported to change in neuronal development (7, 30–32) (Figure 3b). A gene ontology enrichment analysis confirmed that the set of genes that harbor the exons identified by Psix is enriched for genes associated with neuronal and synaptic development (Supplementary Figure 3).

**Fig. 3.**
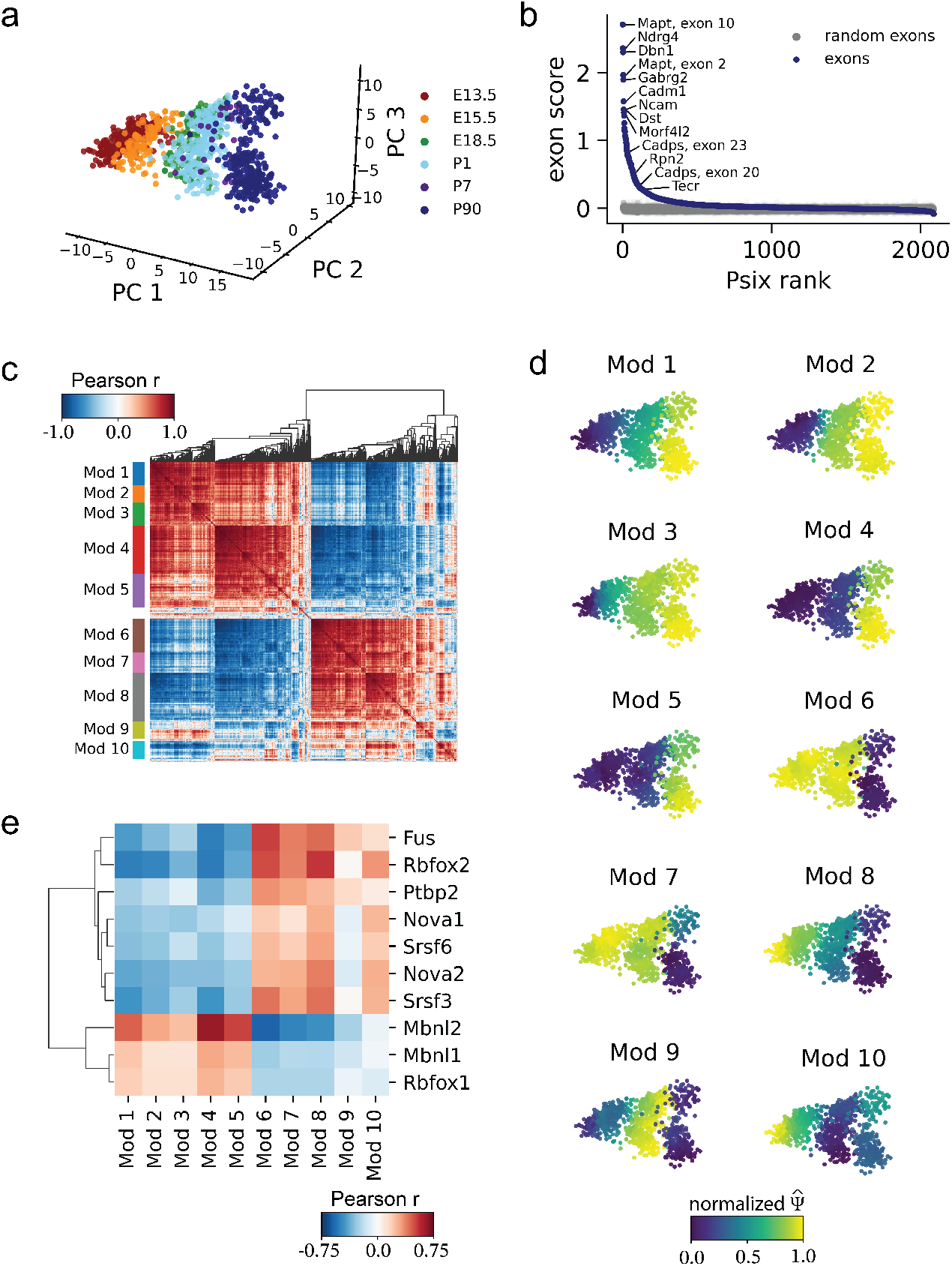
a) Mouse midbrain neurons collected at different stages of development (26), plotted by the first three principle components of normalized gene expression counts. b) Psix scores of cassette exons, compared to the scores of randomized exons. Some exons known to be regulated in neurogenesis are highlighted. c) Correlation map of the neighbor average 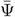 of the cell-state associated exons. Modules were identified with a modification of the UPGMA algorithm. d) Neighbor average normalized 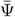 of the exons in each module. e) Correlation of module splicing with gene expression of splicing factors that were enriched for binding in the cell state associated exons identified by Psix. The 10 splicing factors with significant correlation with at least one module (FDR ≤ 0.05, Pearson *r* ≥ 0.25) are shown.

Next we set out to identify co-regulated splicing among the cell-state associated exons. To mitigate the effect that the high variance of individual splicing observations may have on our ability to detect co-regulated exons, we used a neighborhood average Ψ (taking the *k* most similar cells) instead of the individual 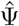. We then clustered these smoothed observations into ten modules of co-spliced exons (Figure 3c). The modules of exons showed distinct patterns of change across brain development, suggesting that they may reflect elements of co-regulation that take place at different developmental phases (Figure 3d). To find potential regulators associated with these modules, we integrated information on splicing factor binding from published CLIP-seq experiments (Table 1) and information on splicing factor expression in single cells. We identified ten splicing factors with enriched binding to the Psix-identified (cell-state associated) exons (hypergeometric test, FDR ≤ 0.05) and whose expression was correlated with the average inclusion rate of exons in at least one of the ten modules (FDR ≤ 0.05, Pearson *r* ≥ 0.25; Figure 3e; Table 1). These splicing factors included proteins from the Nova, Rbfox, Ptbp and Mbnl families, all of which have known roles in regulation of splicing during neuronal development (6, 7, 33–37). Thus, Psix is able to find exons that are coordinately regulated in midbrain development, as well as their potential regulators.

**Table 1.** Mouse CLIP-seq data.

| RBP | Tissue | Binding | Correlation | Source |
| --- | --- | --- | --- | --- |
| CELF1 | C2C12 |  |  | (52) |
| CELF4 | Mouse brain | ✓ |  | (52) |
| ELAVL1 | Mouse brain |  |  | (52) |
| FUS | Mouse brain; mES neurons | ✓ | ✓ | (52) |
| MBNL1 | C2C12 |  |  | (52) |
| MBNL1/2 | Brain, heart, muscle, C2C12 | ✓ | ✓ | (52) |
| MBNL2 | Hippocampus | ✓ | ✓ | (52) |
| NOVA1 | Mouse brain/cortex | ✓ | ✓ | (33, 55) |
| NOVA2 | Mouse cortex | ✓ | ✓ | (55) |
| PABPC1 | MEL |  |  | (52) |
| PTBP1 | C2C12 |  |  | (52) |
| PTBP2 | Mouse embryonic brain | ✓ | ✓ | (35) |
| RBFOX1 | Whole mouse brain | ✓ | ✓ | (52) |
| RBFOX2 | mESC; Whole mouse brain | ✓ | ✓ | (52) |
| RBFOX3 | P19-derived neurons | ✓ |  | (52) |
| RBM3 | MEF |  |  | (52) |
| SRRM4 | N2A cells |  |  | (52) |
| SRSF1 | P19 cells |  |  | (56) |
| SRSF2 | P19 cells |  |  | (56) |
| SRSF3 | P19 cells | ✓ | ✓ | (56) |
| SRSF4 | P19 cells | ✓ |  | (56) |
| SRSF5 | P19 cells | ✓ |  | (56) |
| SRSF6 | P19 cells | ✓ | ✓ | (56) |
| SRSF7 | P19 cells |  |  | (56) |
| TARDBP | Whole mouse brain; Mouse brain | ✓ |  | (52) |
| U2AF2 | N2A cells; Mouse brain | ✓ |  | (52) |

Several splicing modules showed variation within the cells collected at the final time point in the midbrain development timecourse (Figure 3d), and we aimed to leverage the high resolution of single-cell data in understanding this hidden heterogeneity. The neurons collected at the final time point, from adult midbrains 90 days post natal, formed several sub-populations that were separated primarily along the third principal component. The subpopulations express markers of different lineages of midbrain neurons (26) (Supplementary Figure 4). To identify cell-type specific exon usage among these subpopulations, we carried out a separate Psix analysis of just the postnatal day 90 cells (to which we refer as P90 cells). Psix identified 78 exons associated with the variation among neurons in the adult midbrain and grouped them in two modules (Figure 4a). The two modules showed opposite patterns of exon inclusion: module A exons decreased in inclusion along principal component 3, and module B exons increased in inclusion. To trace the emergence of this coordinated pattern during earlier development, we examined the splicing of the same exons at earlier time points (Figure 4a). Similar but weaker correlation patterns were visible in postnatal day 1 (P1) cells, and these patterns were almost absent in embryonic day 13.5 (E13.5) cells, suggesting that these exons responded to differences in splicing factor activity that were present at intermediate stages and became more pronounced with time.

**Fig. 4.**
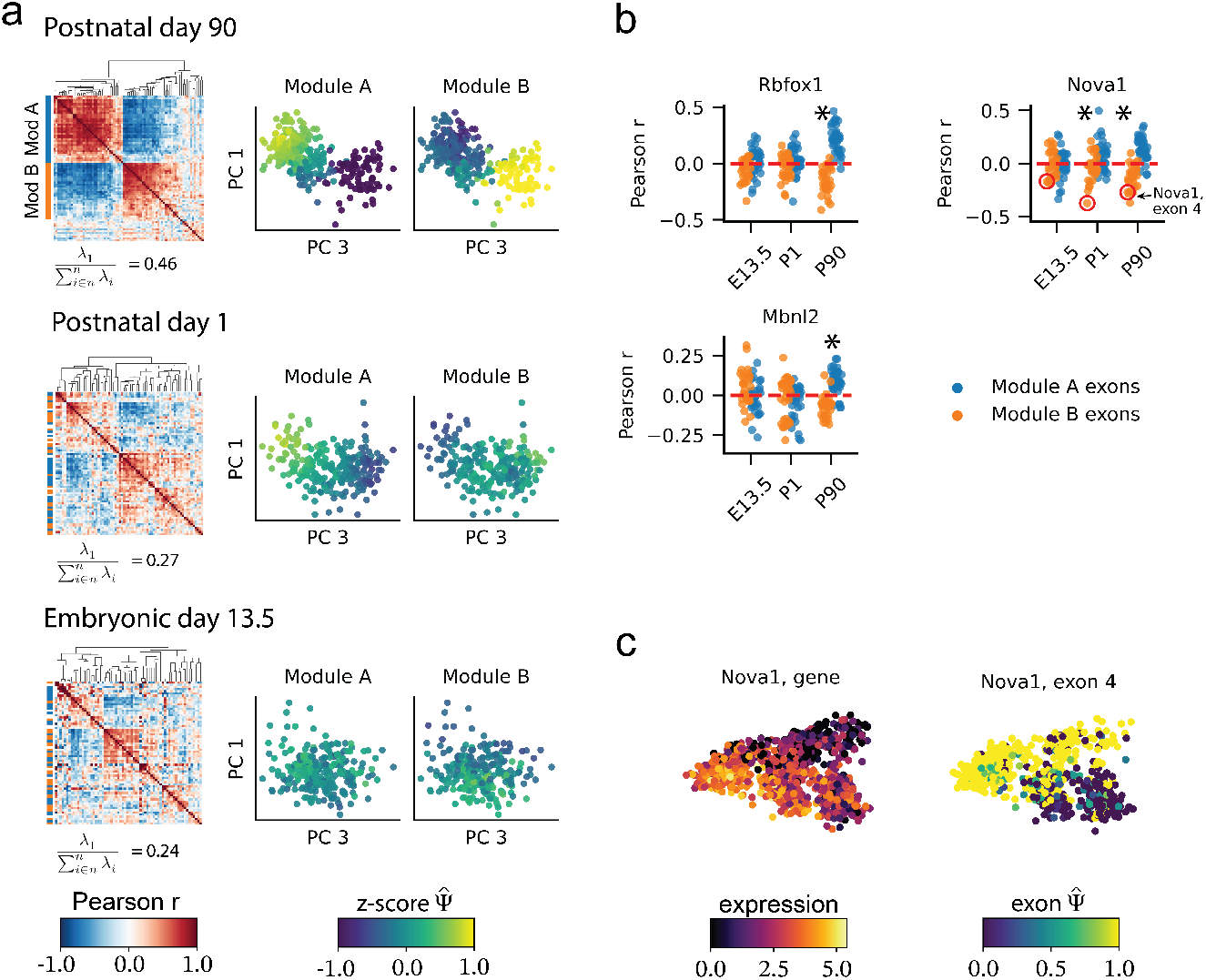
a) Modules of cell-state associated exons identified in postnatal day 90 cells, at different stages of development. Heatmaps show the correlation matrix among the P90 module A and B exons at each time point. As a measure of structure, for each correlation matrix, we show its first eigenvalue divided by the sum of all eigenvalues. Scatterplots show the splicing patterns of the exons of the module in the cells of each time point. b) Correlation of splicing factor expression with the observed 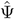 of exons from P90 modules A and B at different stages of development. An asterisk (*) indicates significant difference between the exons of module A and B (Wilcoxon rank test, FDR ≤ 0.05), and a mean difference larger than 0.1 between the two modules. Exon 4 of Nova1 (in module B) is highlighted in red in the correlation plot of Nova1 gene expression. c) Gene expression of Nova1 and splicing of Nova1 exon 4 in mouse midbrain neurons.

To uncover the regulatory mechanisms that control the emergence of these splicing patterns, we first identified seven splicing factors with enriched binding to the Psix-identified, cell-state associated exons in P90 cells. We then computed the correlation between the expression of each splicing factor and the inclusion rate of each exon in modules A and B. Three splicing factors, Nova1, Rbfox1, and Mbnl2, showed a significant difference in their correlation between module A and B (Wilcoxon rank test, FDR ≤ 0.5 and mean difference ≥ 0.1; Figure 4b). Extending the analysis to earlier stages, we found that expression of Nova1 showed a significant difference in correlation with the splicing of the two groups of exons in the P1 cells as well, while the other splicing factors did not show this effect at earlier stages of development. This suggests that Nova1 plays an early role in the emergence of key splicing differences between cell types. In keeping with this role, we observed that one of the exons most highly correlated to Nova1 expression was exon 4 of the Nova1 gene itself, a cassette exon that codes a phosphorylation target domain. In an autoregulatory process, Nova1 protein binding to this exon suppresses its inclusion (38). Consistent with this previous knowledge, we found that expression of Nova1 is strongly anti-correlated with inclusion of exon 4 (Figure 4b). The strength of this correlation is stronger in later stages of development, as one lineage of neurons maintains high Nova1 expression and low exon 4 inclusion, while the other keeps low Nova1 expression and high exon 4 inclusion (Figure 4c). Our results show that the regulatory dynamics of splicing factors and their target exons change during the process of neuron maturation, and consolidate as neurons commit to the distinct midbrain cell lineages. Our analysis exemplifies how single-cell data allows us to study alternative splicing patterns in neurogenesis at high resolution.

We have demonstrated that our probabilistic method, Psix, can find cassette exons that vary continuously across cell state, without mistaking gene expression variance for changes in splicing. Low sequencing coverage has previously limited the study of splicing in single cells (1) and hindered the potential of single cell sequencing to define the identity of individual cells. Psix solves this limitation by combining cell identity information from the transcriptome space with probabilistic modeling that accounts for the distortions of low data recovery. The method could be generalized to use other similarity metrics for single cells, such as spatial positions (39, 40) (to capture spatially-regulated splicing events) or single-cell lineages (41, 42) (to capture heritable splicing events). We have also shown that Psix can identify groups of exons with similar patterns of biological variation, indicative of potential splicing co-regulation. Our methods set the groundwork for further discovery of splicing regulatory networks, taking full advantage of the resolution of single cell methods.

## ACKNOWLEDGEMENTS

We thank Carmelle Catamura for enabling tests of Psix on different data sources and analysis pipelines. C.F.B. was supported by the UC MEXUS-CONACYT doctoral fellowship.

## Methods

### Code and data availability

Psix is available as a Python module at https://github.com/lareaulab/Psix

The analysis of simulations and publicly available data is documented at https://github.com/lareaulab/analysis_psix

The Tiklova (26), Chen (43), and Song (19) datasets were downloaded from accessions GSE116138, GSE74155, and GSE85908, respectively.

### Conceptual overview

To determine if a splicing event is associated with cell identity or cell state, we observe how its Ψ varies in a low-dimensional manifold. If a splicing event is informative about a cell’s identity, we expect that similar cells will have closer Ψ than cells with different identities across the similarity metric.

For a population of *n* single cells, and for *m* alternatively spliced exons, Psix utilizes information from three matrices: one (*n, m*) matrix of 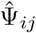 observations for each exon *i* in each cell *j* (obtained from reads covering splice junctions); another (*n, m*) matrix of the estimated number of captured mRNA molecules that cover the splice sites of each exon in each cell; and a low dimensional representations of the cells (Figure 1b). Psix estimates the likelihood of the 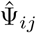 observation of an exon in each cell given the local average Ψ of its closest (k=100) neighbors (based on the low-dimensional representation), and the number of captured molecules *x*_*ij*_. It then contrasts this likelihood with the likelihood of the same observations, given the global average Ψ of all cells. If the likelihood of the observations given the local average is significantly higher than the likelihood given the global average, the exon is considered to be informative of the biological state.

Psix obtains 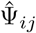 from splicing junction reads of eachanalyzed cassette exon. Because smart-seq2 and similar methods do not directly report the number of mRNAs recovered per gene, for smart-seq2 data Psix estimates the number of mRNAs, *x*_*ij*_, from transcripts per million (TPM) counts for each gene, with a modification of the Census normalization (44) as described in (1). The similarity matrix is estimated from the Euclidean distance in a manifold that summarizes the gene expression space of the single-cell populations. Here, we use the space spanned by the top principal components. Psix could, in principle, use other similarity metrics for single cells not based on gene expression, such as spatial positions.

### The Psix model

Psix is built over the likelihood of 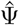 observations based on underlying Ψ and *r* captured mRNA molecules with a capture efficiency *c* that we developed in a previous study (1):

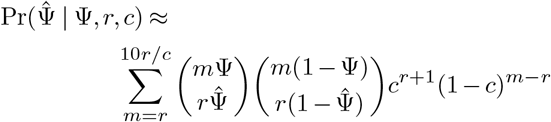

where *m* is the number of molecules present in the cell before sequencing.

Using this likelihood equation, we can test the likelihood of an observation given its underlying splicing context. We compare the likelihood of the observations given that the exon is cell-state associated, vs the likelihood of the observations given that the exon is cell-state independent.

#### Model 1: cell-state associated exons

For an exon *j* and for every cell *i*, we estimate the likelihood of the observed exon inclusion 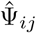, under Model 1: a model in which exon *j* is cell-state associated. First, given a low-dimensional manifold (e.g., the first few principal components of the normalized gene expression), for each cell *i* we define:

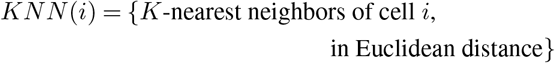

By default, we set *K* = 100 (see Neighbor size selection subsection for details). We also define the sets of cells:

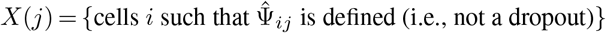

and

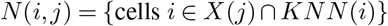

We calculate the neighbor average Ψ for exon *j* in cell *i* as follows:

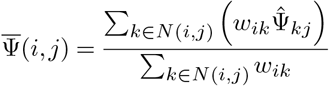

where *K*_*i*_ is the set of k-nearest neighbors of cell *i* (not including itself), and *w*_*ik*_ is a similarity score between cells *i* and *k* in a cell-cell metric, defined from the low-dimensional manifold as follows:

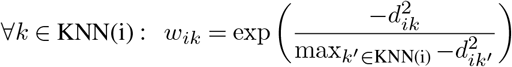

where *d*_*ik*_ corresponds to the Euclidean distance between cells *i* and *k* in the low-dimensional manifold.

We estimate the probability of the observation 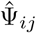 given Model 1 as follows:

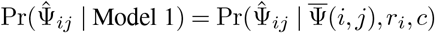

where *r*_*i*_ is the number of captured mRNA molecules that are informative for exon *j* in cell *i*. In rare instances in which a cell’s 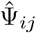 is 0 and the neighbor average 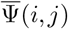 is 1 (or vice-versa), 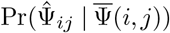 will be equal to 0. To mitigate the impact of these edge cases, we cap this probability to a minimum of 0.01.

#### Model 0: cell-state independent exons

For an exon *j* and for every cell *i*, we estimate the likelihood of the observed exon inclusion 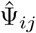 under Model 0, a model in which exon *j* is cell-state independent:

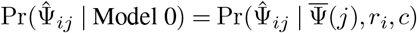

where 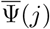 is the unweighted average 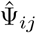 of all cells *i* in *X*(*j*).

#### The Psix score

To determine if an exon is cell-state associated, we compare the likelihood of Model 1 vs Model 0. We define the Psix score as:

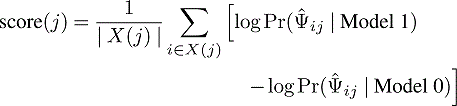

The probability of deviation of these scores from a null distribution is derived empirically, as described below.

### Identifying modules of co-spliced exons

We are interested in identifying exons with similar patterns of cell-state association, since they could potentially share regulatory splicing factors. Due to the high technical noise in single-cell splicing observations, clustering of exons is difficult in the raw data. Instead, for each pair of exons, we obtain the Pearson correlations of the k-nearest neighbors averages. We cluster the exons in a bottom-up procedure similar to that described by DeTomaso & Yosef (40): we merge iteratively the two exons or modules that have the highest Pearson correlation. Then we update the correlation score of the module with all other exons and modules using the UPGMA approach. When a module hits a minimum of 30 exons, we assign a label to it. We stop clustering when the maximal correlation score is lower than 0.3, and return the labeled modules. Exons that do not belong to a labeled module are returned as “unassigned”. The minimum number of exons per module and the minimum correlation score can be defined by the user.

### Practical considerations for the application of the Psix model

#### Estimating captured mRNA molecules in smart-seq2 datasets

In practical usage, non-UMI based scRNA-seq methods such as smart-seq2 do not directly quantify the number of captured mRNA molecules of a gene per cell. As a result, it is difficult to estimate the parameter *r*_*i*_. To approximate *r*, we use a modification of the Census normalization (44) as reported previously:

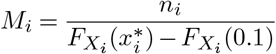

where 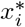 is the mode of the distribution of log-transformed gene TPM counts in cell *i*, 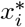 is the argmax of a Gaussian kernel density fit to the log TPM distribution, *n*_*i*_ is the number of genes in cell *i* with a TPM between 0.1 and 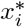, and 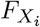 is the cumulative distribution of TPM counts in cell *i*. As in Qiu et al., we assume that genes with TPM below 0.1 are not represented by any mRNA molecules.

#### Low-dimensional manifold

To determine if an exon is informative of a cell’s identity, we need a low-dimensional manifold metric that describes the similarity between individual cells. Due to hurdles such as noise, sparsity, and the curse of dimensionality, it is common practice to define this metric as a Euclidean distance in an interpretable low-dimensional manifold representation of the normalized gene expression profiles. In general the approach to normalization and dimensionality reduction would depend on the size of the dataset analyzed. For the Tiklova (26), Chen (43) and Song (19) datasets, we normalized the TPM counts per gene using SCONE (24) due to the relatively small number of cells, and performed dimensionality reduction with PCA over the top 1000 variable (using the fano factor) genes. In the Chen and Song datasets, the first two principal components captured most of the variance, and thus we used these components as the low-dimensional manifold. For the larger Tiklova dataset, we used the first three principal components.

#### Neighborhood size selection

Our simulation analysis showed that Psix performance was optimal when the size of the neighborhood used for estimating neighbor average 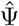 is around ~10% of the total number of cells in the dataset (Supplementary Figure 2a,b).

#### Capture efficiency in single-cell datasets

We set a capture efficiency of 0.1, as supported by single-cell studies (44–47). The choice of capture efficiency has very little effect on Psix scores (Supplementary Figure 2c).

#### Empirical p-value estimation

Psix compares the probability of two models. However, the null model (Model 0) is not embedded into the alternative model (Model 1). For this reason, for each exon *j* we obtained an empirical *p*-value by randomizing the 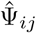 observations of all cells *i* and their respective mRNA counts, and obtaining a random distribution of Psix scores. We observed that the mean mRNA counts and the total 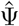 variance had a slight effect on the random distribution of the Psix scores. For this reason, we divided the exons into bins according to their ranks in mRNA counts and 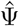 variance (by default, 5 bins for each parameter, totalling 25 bins). For each bin, we randomized the exons of the bin, and for each exon we estimated an empirical *p*-value as follows:

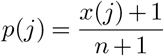

where *x*(*j*) is the number of random exons in the bin to which the exon *j* belongs that have a higher Psix score than the non-randomized exon, and *n* is the total number of randomized exons in the bin (2000 by default). By default, Psix uses the Benjamini-Hochberg procedure to correct for multiple tests.

#### Processing of smart-seq2 datasets

For the mouse mid-brain development dataset (26) we aligned the reads using STAR to the mouse genome (annotation mm10) with added sequences for the Illumina ERCC spike-ins and the enhanced green fluorescent protein (eGFP) gene sequence. TPM was estimated using RSEM. After quality control and removing a substantial number of outlier cells expressing glial or microglial gene markers, we were left with a total of 1067 cells that we used for downstream analysis. For dimensionality reduction, we first selected genes that are expressed in the dataset (average normalized counts ≥ 0.1), and selected the top 1000 genes with high Fano factor. We then applied PCA on the resulting genes, which revealed a clear trajectory of neuron differentiation.

Cassette exons were identified from the mm10 mouse genomic annotation. We estimated the observed 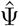for each cassette exon *j* in each cell *i* as:

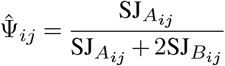

where 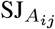 are the total RNA-seq reads that cover the two splice junctions that support exon inclusion of the exon. In turn, 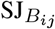 are the total RNA-seq reads that cover the splice junction that support exon exclusion.

We selected 2087 cassette exons that had observations in at least 25% of cells (i.e., a maximum dropout rate of 75%). We applied Psix over these exons using a neighborhood size of 100 cells and a capture efficiency rate of 0.1. After scoring these exons with Psix, Gene Ontology analysis was performed using the Panther Enrichment Test (48) on the Psix score of the exon. For genes with more than one exon, we used the highest scoring exon’s Psix score.

The Chen (43) and Song (19) datasets were processed as we described in our previous work (1). These datasets are smaller than the Tiklova dataset. A neighborhood of size 30 was used in both cases.

### Comparison of Psix with other single-cell splicing methods

We compared the performance of Psix against several approaches that have been proposed for addressing single-cell splicing.

#### Kruskal-Wallis test

The Kruskal-Wallis test is a non-parametric version of ANOVA. It has been previously used to identify if the median 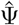 of an exon is different between predefined clusters of cells (1, 21). We applied this test for all the exons we evaluated with Psix in all datasets. We clustered the cells according to the labels provided by the authors of each dataset.

#### Geary’s C

Geary’s C is a statistic that measures the spatial autocorrelation of a variable. It has been used for computing the autocorrelation of gene signatures with cell identity (49), and we previously used an adaptation of this test for finding autocorrelation of exon observations (1).

For each exon, we normalize the observed 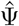 as follows:

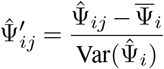

where 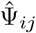 is the observed splicing of exon *j* in cell *i*. 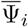 is the mean 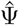 of all the observed exons in cell *i*, and 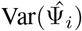 is the variance of the observed 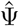 of all exons in cell *i*.

We calculate a variant of the Geary’s C statistic of each exon *j* as:

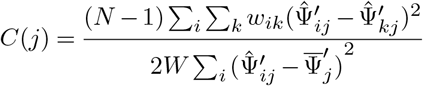

where *N* is the number of cells in the dataset; 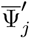 is the average 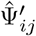 of exon *j* across all cells; *w*_*i,k*_ is the weight between cell *i* and cell *k* in the cell-cell metric, and *W* is the sum of all *w*_*i,k*_. We use the same similarity metric that we used in Psix.

In order to make the statistic more intuitive, in which a positive score close to 1 indicates high autocorrelation, we transform our statistic as:

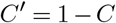

#### BRIE2

BRIE2 is a computational tool that regresses single-cell splicing data against cell-level features, and can detect differential splicing between groups of cells.

We applied BRIE2 directly to the STAR-aligned BAM files (50) of the Tiklova, Chen and Song datasets. We used the author-provided labels for each dataset.

#### Alternative splicing in bulk RNA-seq data

We compared each method’s ability to detect alternative splicing events identified in bulk RNA-seq datasets using rMATS 3.2.5 (27). We compared the Tiklova dataset (26) with a bulk mouse brain development dataset (7); the Chen dataset (43) with a bulk dataset of mouse embryonic stem cells induced to neurogenesis (51); and the Song dataset with the bulk RNA-seq data included in the same study (19).

### RNA binding protein analysis

CLIP-seq data was obtained from multiple public sources (Supplementary Table 1). Some datasets were downloaded from the CLIPdb dataset (52), while others were obtained from their respective studies depending on availability. For the SR protein iCLIP-seq data that was not already pre-processed by CLIPdb, we used Piranha version 1.2.1 (53) to call peaks from CLIP-tags. Peaks from different datasets of the same RNA binding protein (RBP) were combined into a single file per RBP. We next reported the overlap of the RBPs into four regions per cassette exon:

- E1 region: 100 nt upstream and *i*_1_ nt downstream from the splice junction of the upstream flanking exon. Where *i*_1_ is the minimum of 100 and half the distance between the upstream flanking exon and the cassette exon.
- S1 region: *i*_1_ nt upstream and *e* nt downstream from the 5’ splice junction of the cassette exon. Where *e* is the minimum of 100 and half the distance between the two splice junctions of the cassette exon.
- S2 region: *e* nt upstream and *i*_2_ nt downstream from the 3’ splice junction of the cassette exon. Where *i*_2_ is the minimum of 100 and half the distance between the cassette exon and the downstream flanking exon.
- E2 region: *i*_2_ nt upstream and 100 nt downstream from the splice junction of the downstream flanking exon.

We consider that a splicing factor binds to the exon if it binds to any of the four regions. For the Tiklova dataset, we tested the binding enrichment of each RBP independently in the 798 cell-state associated exons identified by Psix, using the hypergeometric test, with the 2,087 tested exons as the background. We applied the Benjamini-Hochberg procedure to correct for multiple testing for all the RBPs. We used the same approach to test for enrichment in the 78 cell-state associated exons in the P90 cells versus the background of 2,115 tested exons.

#### Analysis of postnatal day 90 modules

We implemented Psix on the 290 cells collected from the postnatal day 90 midbrain (P90 cells), using the transcriptomic space in Figure 3a. Due to the smaller sample size (compared with the entire dataset), we used a neighborhood of 30 cells (~10% of the total number of cells), and a minimum correlation for neighbor joining of 0.1. This resulted in 78 cell-state associated exons, and two modules.

In order to find the potential regulators that control the formation of these patterns during neurogenesis, we identified the RNA binding proteins that are significantly enriched in binding to the 78 exons in these modules using CLIP-seq data (hypergeometric test, FDR ≤ 0.05). We tested for enrichment of binding to any of the four regions of the exon previously described.

We ran a Pearson correlation test between the normalized expression (SCONE normalized TPM counts) of the enriched splicing factors and the observed 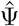 of the exons in each module in P90 cells. We tested for significance in the difference in distribution of Pearson correlation scores using the Wilcoxon rank test, and correcting for multiple testing with the Benjamini-Horchberg correction. We repeated these observations with cells from the postnatal day 1 (P1 cells), and embryonic day 13.5 (E13.5) stages of development.

### Simulations of alternative splicing in single cells

We simulate alternative splicing of exons in single cells. This consists of three main steps: First, we simulate gene expression profiles in a population of single-cells sampled from a continuous trajectory; with either a single branch (Figure 2a) or multiple (Supplementary Figure 1a, b). Second, we simulate alternative splicing in the exons of this population (accounting for one exon per gene), where in some exons the inclusion rates are associated with the trajectory and in some the rates are random. These first two parts serve to simulate the true molecular content of the cells. Third, we simulate the measurement process (e.g., mRNA capture, amplification, sequencing), which results in noisy and sparse data, such as limited availability of splice junction reads.

#### Step 1: Simulating gene expression in a single-cell population

We used SymSim (25) to simulate the expression of 5000 genes in a continuous population of 1000 single cells with default parameters. To simulate a single developmental trajectory, we simulated single cells across a phylogenetic tree with two branches with SymSim. To simulate a single lineage, we set the endpoint of one branch as the starting point of the lineage, and the endpoint of the second branch as the end of the lineage. Each cell’s relative position in the tree corresponded to its position in the developmental trajectory, starting from the tip of one branch to the other. We normalized the relative positions of the lineage to span from 0 (first cell) to 100 (last cell).

#### Step 2: Simulating splicing in single cells

We simulate splicing in the single cells as three sub-steps. First, we model the expected Ψ as an impulse function, which was shown by us and others to fit well with empirical time course data (54). For each exon, we simulate a “Platonic” Ψ that will be dependent on the cell’s position in the lineage:

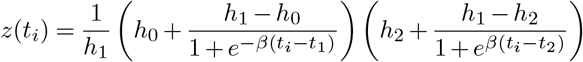

where *z*(*t*_*i*_) = logit(Ψ(*t*_*i*_)). *t*_*i*_ ∈ [0, 100] is the position of a given cell in the simulated lineage. *h*_0_, *h*_1_ and *h*_2_ are amplitude parameters of the curve (first, second and third plateau), *t*_1_ and *t*_2_ are the state transition times; i.e., when the inflection points of the impulse happen. *β* is a slope parameter that determines how steep the impulse is.

We sample the impulse parameters from the following probabilistic distributions:

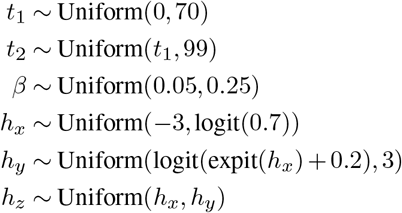

To simulate different impulse shapes, we set [*h*_0_, *h*_1_, *h*_2_] as a permutation of [*h*_*x*_, *h*_*y*_, *h*_*z*_]. To simulate exons that do not change across the lineage, we randomly select a value *x* from [*h*_*x*_, *h*_*y*_, *h*_*z*_] and set *h*_0_ = *h*_1_ = *h*_2_ = *x* to specify a horizontal line.

We add noise to simulate the biological variability that affects the true splicing profile of a cell in a manner independent of the lineage:

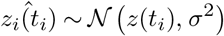

where *z*_*i*_(*t*_*i*_) is the logit transformation of the true Ψ of cell *i*. *t*_*i*_ is the temporal stage of cell *i*; i.e., its position in the continuous trajectory (lineage). *σ*^2^ is a variance parameter that is specific of each exon. For each exon, we randomly sample *σ*^2^ ~ Uniform(0.5, 1). Finally, we simulate splicing of the mRNA molecules of each gene as a stochastic process as follows:

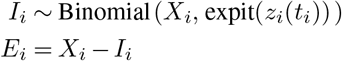

where *I*_*i*_ is the number of mRNA molecules of the gene in cell *i* that include the alternatively spliced exon. *E*_*i*_ is the number of mRNA molecules that exclude the exon. *X*_*i*_ is the total number of mRNA molecules of the gene in cell *i*. We merge *E* and *I* into a single matrix of mRNA molecules *M*, in which each isoform corresponds to an independent row.

#### Step 3: Sequencing and splice junction coverage

Once we have simulated the expression and splicing of each gene, we simulate the sequencing of the mRNA molecules. For this, we first assign a transcript length to each molecule as follows: For each gene *g* with isoforms *g*_*E*_ (excludes the exon) and *g*_*I*_ (includes the exon), we randomly sample without replacement 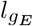 as the length of *g*_*E*_ from SymSim’s transcript length database. We then sample without replacement 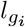, the length of the alternative exon of *g*, from a database of lengths of cassette exons in the human genome. Finally, we set the length of *g*_*I*_ as 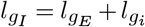.

We use SymSim’s `True2ObservedCounts` function to simulate non-UMI sequencing of the mRNA molecules in *M* and lengths 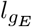 and 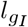. We used the following parameters: mean capture efficiency: 0.1, capture efficiency standard deviation: 0.05, depth sequencing mean: 1e5, depth sequencing standard deviation: 1e4. To simulate a dataset with bad capture efficiency, we set mean capture efficiency: 0.05, capture efficiency standard deviation: 0.02; and for a dataset with very poor capture efficiency we set mean capture efficiency: 0.01, capture efficiency standard deviation: 0.01.

Finally, we simulate the splice junction coverage of isoforms as follows:

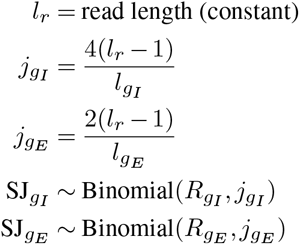

where *l*_*r*_ corresponds to the constant read length from the sequencing process (set as default to 50). 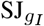 and 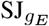 are the number of reads that cover informative splice junctions for isoforms *g*_*I*_ and *g*_*E*_ respectively; and 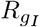 and 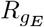 are the total number of reads simulated to map to isoforms *g*_*I*_ and *g*_*E*_ respectively.

For each cell, the observed 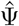of each exon is calculated as:

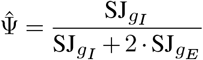

#### Simulation of a branching lineage of single cells

We simulated a diverging lineage of single cells, using SymSim with the following three-branched phylogenetic tree (Supplementary Figure 1):

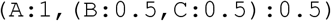

We divided the single-cell population into three lineages:

- Lineage 1: Root to A
- Lineage 2: Root to B
- Lineage 3: B-C splitting point to C

We simulated the “Platonic” Ψ of each lineage as follows:

1. For Lineage 1, sample *h*_0_, *h*_1_, *t*_1_ and *β* as shown for the single-lineage impulse parameters. With a probability of 0.25, simulate differential splicing with a sigmoid function as follows:

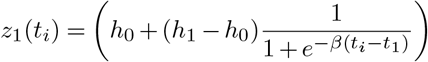

Else, we fit a flat line *z*_1_(*t*_*i*_) = *h*_0_.
2. For Lineage 2, we use the same *h*_0_ as in Lineage 1, and randomly sample the other parameters. We simulate differential splicing with a sigmoid *z*_2_ with a probability of 0.25, or fit a flat line *z*_2_(*t*_*i*_) = *h*_0_ otherwise.
3. For Lineage 3, we set *h*_0_ = *z*_2_(*t*_*BC*_), where *z*_2_ is the function (sigmoid or flat line) used for Lineage 2, and *t_BC_* is the timepoint at which Lineage 2 and Lineage 3 split. We sample the other parameters randomly. We simulate splicing with a sigmoid *z*_3_ with a probability of 0.25, or fit a flat line *z*_3_(*t*_*i*_) = *h*_0_ otherwise.

Each simulated gene is marked as differentially spliced if the splicing of at least one lineage was simulated with a sigmoid. Otherwise the gene is marked as non-differentially spliced. Once we’ve simulated the “Platonic” Ψ of each gene, we repeat the simulation steps for the single lineage simulations.

---

**Supplementary Figure 1.**
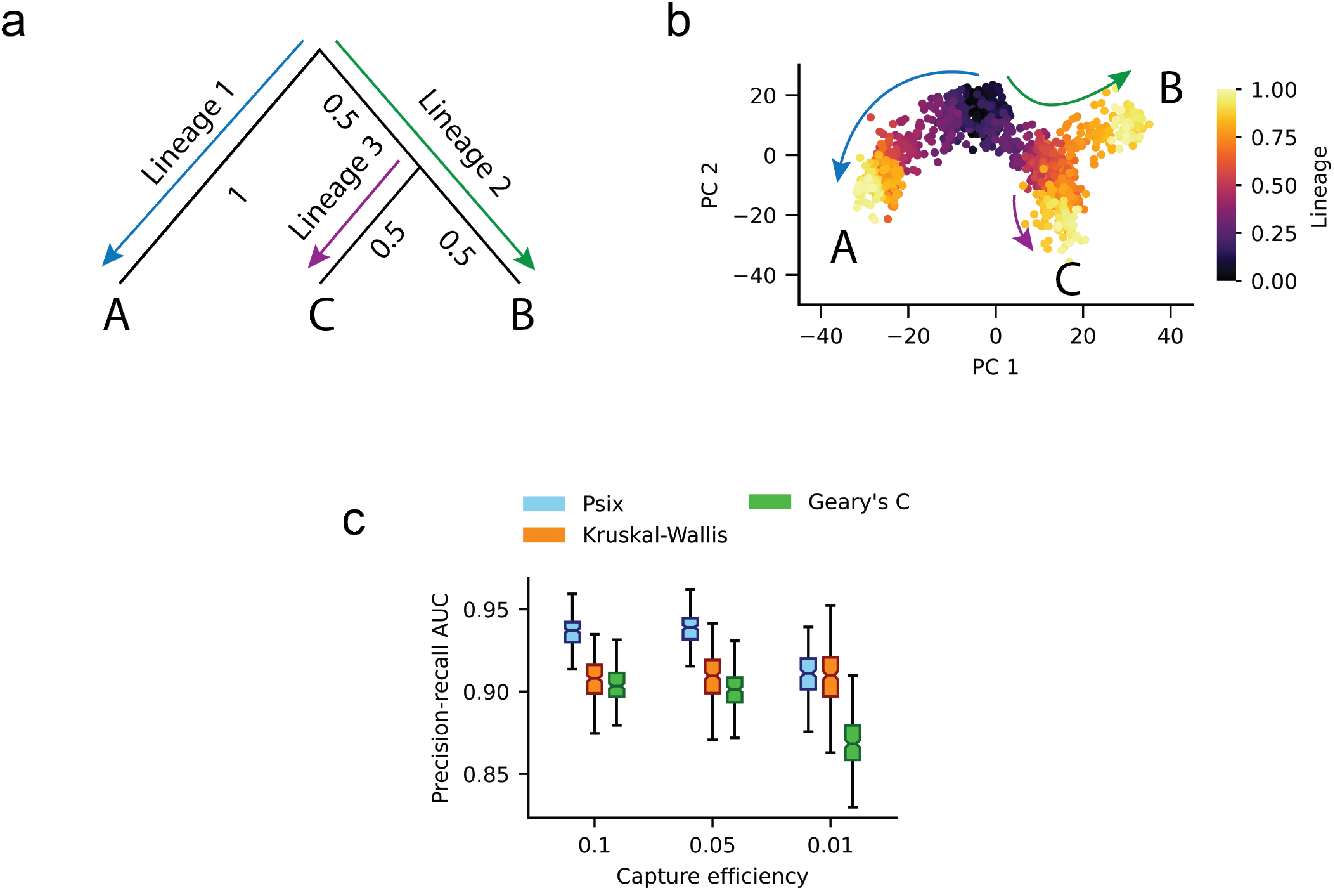
Simulation of alternative splicing in a complex single-cell population. a) Phylogenetic tree used as input for SymSim. b) Principal component analysis of the transcriptome landscape of the simulated population. c) Area under the precision-recall curve of Psix, the Kruskal-Wallis test, and Geary’s C, identifying exons with a | ΔΨ | ≥ 0.2 under different capture efficiencies.

**Supplementary Figure 2.**
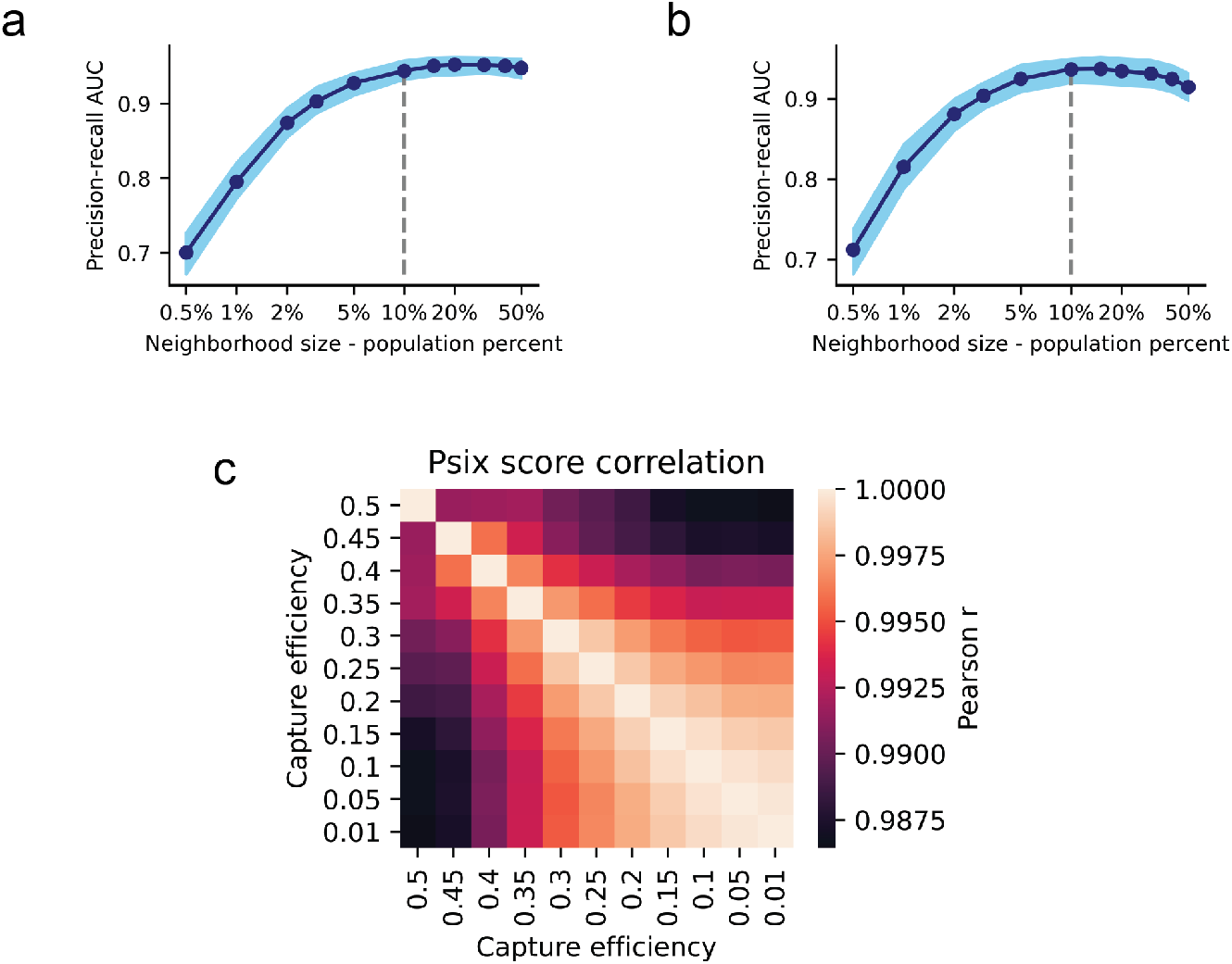
a,b) Effect of neighborhood size on Psix performance. Area under the precision-recall curve of Psix on simulated data with different *k* sizes. Blue area represents 0.05 to 0.95 quantiles. Data is shown from a) the single lineage simulation, and from b) the three lineages simulation. c) Pearson correlation of Psix score in the Tiklova dataset (26) using different capture efficiency rates in the model.

**Supplementary Figure 3.**
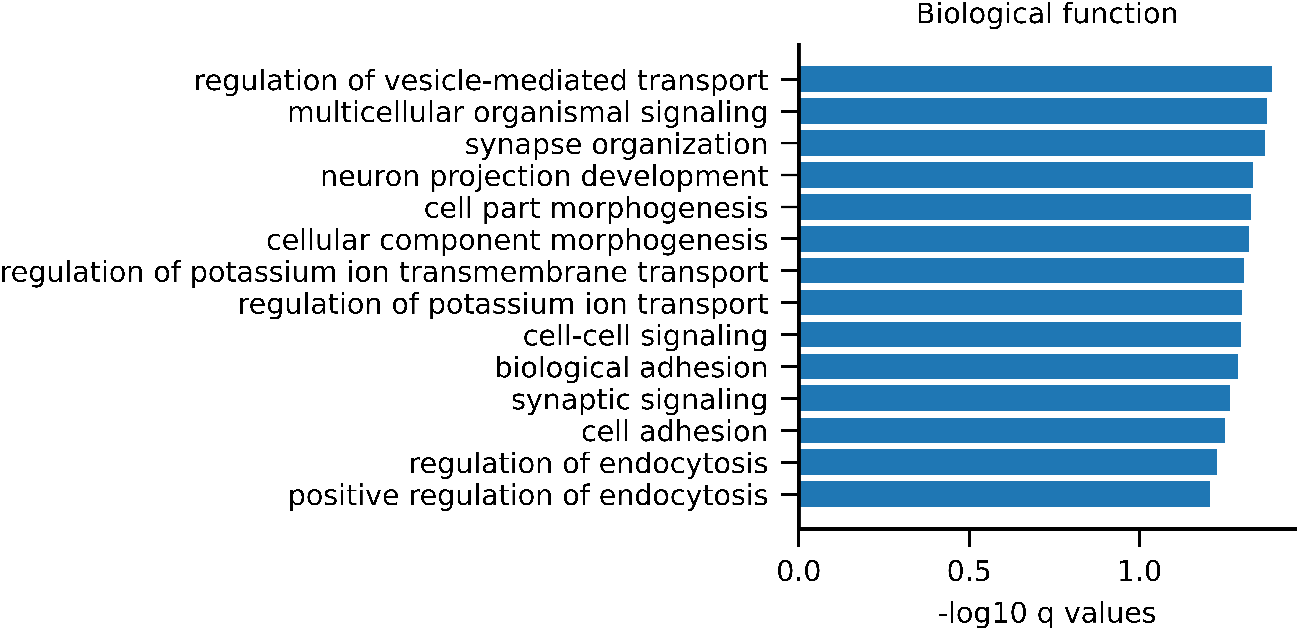
Gene Ontology enrichment analysis of the Psix score in the Tiklova dataset (biological function terms).

**Supplementary Figure 4.**
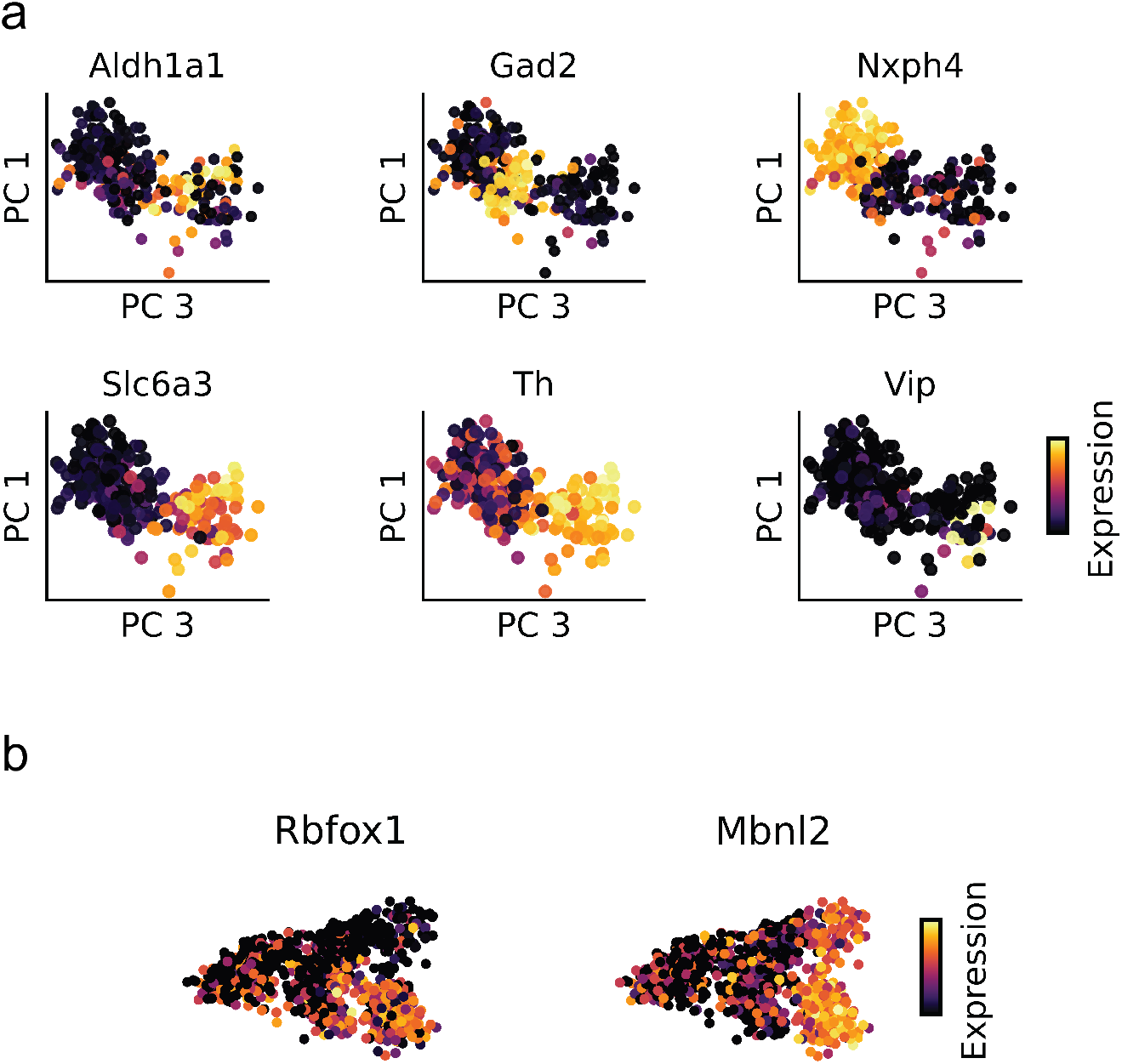
a) Principal component 3 is associated with neuron diversity in postnatal day 90 midbrain neurons. Expression of genes characteristic of midbrain neuron subtypes. b) Gene expression of Rbfox1and Mbnl2 in midbrain dopamine neurons during development.

## Bibliography

1. Carlos F Buen Abad Najar, Nir Yosef, and Liana F Lareau. Coverage-dependent bias creates the appearance of binary splicing in single cells. eLife, 9:e54603, 2020.

2. Allon Wagner, Aviv Regev, and Nir Yosef. Revealing the vectors of cellular identity with single-cell genomics. Nature Biotechnology, 34(11):1145–1160, 2016.

3. Amos Tanay and Aviv Regev. Scaling single-cell genomics from phenomenology to mechanism. Nature, 541(7637):331–338, 2017.

4. Eric T Wang, Rickard Sandberg, Shujun Luo, Irina Khrebtukova, Lu Zhang, Christine Mayr, Stephen F Kingsmore, Gary P Schroth, and Christopher B Burge. Alternative isoform regulation in human tissue transcriptomes. Nature, 456(7221):470–476, 2008.

5. Francisco E Baralle and Jimena Giudice. Alternative splicing as a regulator of development and tissue identity. Nature Reviews Molecular Cell Biology, 18(7):437–451, 2017.

6. Bushra Raj and Blencowe Benjamin J. Alternative Splicing in the Mammalian Nervous System: Recent Insights into Mechanisms and Functional Roles. Neuron, 87(1):14–27, 2015.

7. Sebastien M Weyn-Vanhentenryck, Huijuan Feng, Dmytro Ustianenko, Rachel Duffié, Qinghong Yan, Martin Jacko, Jose C Martinez, Marianne Goodwin, Xuegong Zhang, Ulrich Hengst, Stavros Lomvardas, Maurice S Swanson, and Chaolin Zhang. Precise temporal regulation of alternative splicing during neural development. Nature Communications, 9(1): 2189–17, 2018.

8. Kenichi Yoshida, Masashi Sanada, Yuichi Shiraishi, Daniel Nowak, Yasunobu Nagata, Ryo Yamamoto, Yusuke Sato, Aiko Sato-Otsubo, Ayana Kon, Masao Nagasaki, George Chalkidis, Yutaka Suzuki, Masashi Shiosaka, Ryoichiro Kawahata, Tomoyuki Yamaguchi, Makoto Otsu, Naoshi Obara, Mamiko Sakata-Yanagimoto, Ken Ishiyama, Hiraku Mori, Florian Nolte, Wolf-Karsten Hofmann, Shuichi Miyawaki, Sumio Sugano, Claudia Haferlach, H. Phillip Koeffler, Lee-Yung Shih, Torsten Haferlach, Shigeru Chiba, Hiromitsu Nakauchi, Satoru Miyano, and Seishi Ogawa. Frequent pathway mutations of splicing machinery in myelodysplasia. Nature, 478(7367):64–69, 2011.

9. Manuel Irimia, Robert J Weatheritt, Jonathan D Ellis, Neelroop N Parikshak, Thomas Gonatopoulos-Pournatzis, Mariana Babor, Mathieu Quesnel-Vallières, Javier Tapial, Bushra Raj, Dave O’Hanlon, Miriam Barrios-Rodiles, Michael J Sternberg, Sabine P Cordes, Frederick P Roth, Jeffrey L Wrana, Daniel H Geschwind, and Benjamin J Blencowe. A highly conserved program of neuronal microexons is misregulated in autistic brains. Cell, 159(7): 1511–1523, 2014.

10. Héctor Climente-González, Eduard Porta-Pardo, Adam Godzik, and Eduardo Eyras. The functional impact of alternative splicing in cancer. Cell Reports, 20(9):2215–2226, 2017.

11. Neelroop N. Parikshak, Vivek Swarup, T. Grant Belgard, Manuel Irimia, Gokul Ramaswami, Michael J. Gandal, Christopher Hartl, Virpi Leppa, Luis de la Torre Ubieta, Jerry Huang, Jennifer K. Lowe, Benjamin J. Blencowe, Steve Horvath, and Daniel H. Geschwind. Genome-wide changes in lncrna, splicing, and regional gene expression patterns in autism. Nature, 540(7633):423–427, 2016.

12. Davide Risso, Fanny Perraudeau, Svetlana Gribkova, Sandrine Dudoit, and Jean-Philippe Vert. A general and flexible method for signal extraction from single-cell RNA-seq data. Nature Communications, 9(284), 2018.

13. Romain Lopez, Jeffrey Regier, Michael B Cole, Michael I Jordan, and Nir Yosef. Deep generative modeling for single-cell transcriptomics. Nature Methods, 15(12):1053–1058, 2018.

14. Gökcen Eraslan, Lukas M Simon, Maria Mircea, Nikola S Mueller, and Fabian J Theis. Single-cell RNA-seq denoising using a deep count autoencoder. Nature Communications, 10(390), 2019.

15. Tim Stuart, Andrew Butler, Paul Hoffman, Christoph Hafemeister, Papalexi, Efthymia, William M Mauck, Yuhan Hao, Marlon Stoeckius, Peter Smibert, and Rahul Satija. Comprehensive Integration of Single-Cell Data. Cell, 177(7):1888–1902, 2019.

16. Joshua D Welch, Velina Kozareva, Ashley Ferreira, Charles R Vanderburg, Carly A Martin, and Evan Z Macosko. Single-Cell Multi-omic Integration Compares and Contrasts Features of Brain Cell Identity. Cell, 177(7):1873–1887, 2019.

17. Jennifer Westoby, Pavel Artemov, Martin Hemberg, and Anne Ferguson-Smith. Obstacles to detecting isoforms using full-length scRNA-seq data. Genome Biology, 21(74), 2020.

18. Alex K Shalek, Rahul Satija, Xian Adiconis, Rona S Gertner, Jellert T Gaublomme, Raktima Raychowdhury, Schraga Schwartz, Nir Yosef, Christine Malboeuf, Diana Lu, John J Trombetta, Dave Gennert, Andreas Gnirke, Alon Goren, Nir Hacohen, Joshua Z Levin, Hongkun Park, and Aviv Regev. Single-cell transcriptomics reveals bimodality in expression and splicing in immune cells. Nature, 498(7453):236–240, 2013.

19. Yan Song, Olga B Botvinnik, Michael T Lovci, Boyko Kakaradov, Patrick Liu, Jia L Xu, and Gene W Yeo. Single-Cell Alternative Splicing Analysis with Expedition Reveals Splicing Dynamics during Neuron Differentiation. Molecular Cell, 67(1):148–161.e5, 2017.

20. Joshua D Welch, Yin Hu, and Jan F Prins. Robust detection of alternative splicing in a population of single cells. Nucleic Acids Research, 44(8):e73–e73, 2016.

21. Wei Xiong Wen, Adam J Mead, and Supat Thongjuea. VALERIE: Visual-based inspection of alternative splicing events at single-cell resolution. PLoS Computational Biology, 16(9): e1008195, 2020.

22. Yuanhua Huang and Guido Sanguinetti. BRIE: transcriptome-wide splicing quantification in single cells. Genome Biology, 18(1):123, 2017.

23. Yuanhua Huang and Guido Sanguinetti. Computational identification of splicing phenotypes from single cell transcriptomic experiments. biorxiv, 2020. doi: https://doi.org/10.1101/2020.11.04.368019.

24. Michael B Cole, Davide Risso, Allon Wagner, David DeTomaso, John Ngai, Elizabeth Purdom, Sandrine Dudoit, and Nir Yosef. Performance Assessment and Selection of Normalization Procedures for Single-Cell RNA-Seq. Cell Systems, 8(4):315–328.e8, 2019.

25. Xiuwei Zhang, Chenling Xu, and Nir Yosef. Simulating multiple faceted variability in single cell RNA sequencing. Nature Communications, 10:2611–27, 2019.

26. Katarína Tiklová, Asa K Björklund, Laura Lahti, Alessandro Fiorenzano, Sara Nolbrant, Linda Gillberg, Nikolaos Volakakis, Chika Yokota, Markus M Hilscher, Thomas Hauling, Fredrik Holmström, Eliza Joodmardi, Mats Nilsson, Malin Parmar, and Thomas Perlmann. Single-cell RNA sequencing reveals midbrain dopamine neuron diversity emerging during mouse brain development. Nature Communications, 10(581), 2019.

27. Shihao Shen, Juw Won Park, Zhi-Xiang Lu, Lan Lin, Michael D Henry, Ying Nian Wu, Qing Zhou, and Yi Xing. rMATS: robust and flexible detection of differential alternative splicing from replicate RNA-Seq data. Proceedings of the National Academy of Sciences of the United States of America, 111(51):E5593–601, 2014.

28. Sara Aibar, Carmen Bravo González-Blas, Thomas Moerman, Vân Anh Huynh-Thu, Hana Imrichova, Gert Hulselmans, Florian Rambow, Jean-Christophe Marine, Pierre Geurts, Jan Aerts, Joost van den Oord, Zeynep Kalender Atak, Jasper Wouters, and Stein Aerts. Scenic: single-cell regulatory network inference and clustering. Nature Methods, 14(11):1083–1086, 2017.

29. Hiujuan Feng, Daniel F Moakley, Shuonan Chen, Melissa G McKenzie, Vilas Menon, and Chaolin Zhang. Complexity and graded regulation of neuronal cell-type–specific alternative splicing revealed by single-cell rna sequencing. Proceedings of the National Academy of Sciences of the United States of America, 118(10):e2013056118, 2021.

30. Pascal Y Smith, Charlotte Delay, Johanne Girard, Marie-Amélie Papon, Emmanuel Planel, Nicolas Sargeant, Luc Buée, and Sébastien S Hébert. Microrna-132 loss is associated with tau exon 10 inclusion in progressive supranuclear palsy. Human Molecular Genetics, 20 (20):4016–4024, 2011.

31. Jia Bei Wang and David R Burt. Differential expression of two forms of gabaa receptor gamma 2-subunit in mice. Brain Research Bulletin, 27(5):731–735, 1991.

32. Dina Speidel, Frederique Varoqueaux, Carsten Enk, Mari Nojiri, Ruslan N Grishanin, Thomas FJ Martin, Kay Hoffman, Nils Brose, and Kerstin Reim. A family of ca2+-dependent activator proteins for secretion: comparative analysis of structure, expression, localization, and function. Journal of Biological Chemistry, 278(52):52802–52809, 2003.

33. Chaolin Zhang, Maria A Frias, Aldo Mele, Matteo Ruggiu, Taesun Eom, Christina B Marney, Huidong Wang, Donny D Licatalosi, John J Fak, and Robert B Darnell. Integrative modeling defines the Nova splicing-regulatory network and its combinatorial controls. Science, 329 (5990):439–443, 2010.

34. Konstantinos Charizanis, Kuang-Yung Lee, Ranjan Batra, Marianne Goodwin, Chaolin Zhang, Yuan Yuan, Lily Shiue, Melissa Cline, Marina M. Scotti, Guangbin Xia, Ashok Kumar, Tetsuo Ashizawa, H. Brent Clark, Takashi Kimura, Masanori P. Takahashi, Harutoshi Fujimura, Kenji Jinnai, Hiroo Yoshikawa, Mário Gomes-Pereira, Geneviève Gourdon, Noriaki Sakai, Seiji Nishino, Thomas C. Foster, Manuel Ares, Robert B. Darnell, and Maurice S. Swanson. Muscleblind-like 2-mediated alternative splicing in the developing brain and dysregulation in myotonic dystrophy. Neuron, 75(3):437–450, 2012.

35. Donny D Licatalosi, Masato Yano, John J Fak, Aldo Mele, Sarah E Grabinski, Chaolin Zhang, and Robert B Darnell. Ptbp2 represses adult-specific splicing to regulate the generation of neuronal precursors in the embryonic brain. Genes & Development, 26(14):1626–1642, 2012.

36. Qin Li, Sika Zheng, Areum Han, Chia-Ho Lin, Peter Stoilov, Xiang-Dong Fu, and Douglas L Black. The splicing regulator PTBP2 controls a program of embryonic splicing required for neuronal maturation. eLife, 3:e01201, 2014.

37. Sebastien M Weyn-Vanhentenryck, Aldo Mele, Qinghong Yan, Shuying Sun, Natalie Farny, Zuo Zhang, Chenghai Xue, Margaret Herre, Pamela A Silver, Michael Q Zhang, Adrian R Krainer, Robert B Darnell, and Chaolin Zhang. HITS-CLIP and integrative modeling define the Rbfox splicing-regulatory network linked to brain development and autism. Cell Reports, 6(6):1139–1152, 2014.

38. B Kate Dredge, Giovanni Stefani, Caitlin C. Engelhard, and Robert B Darnell. Nova autoregulation reveals dual functions in neuronal splicing. EMBO Journal, 24(8):1608–1620, 2005.

39. Svensson Valentine, Sarah A Teichmann, and Oliver Stegle. SpatialDE: identification of spatially variable genes. Nature Methods, 15(5):343–346, 2018.

40. David DeTomaso and Nir Yosef. Hotspot identifies informative gene modules across modalities of single-cell genomics. Cell Systems, 12(5):446–456, 2021.

41. Lennart Kester and Alexander van Oudenaarden. Single-cell transcriptomics meets lineage tracing. Cell Stem Cell, 23(2):166–179, 2018.

42. Matthew G Jones, Alex Khodaverdian, Jeffrey J Quinn, Michelle M Chan, Jeffrey A Hussmann, Robert Wang, Chenling Xu, Jonathan S Weissman, and Nir Yosef. Inference of single-cell phylogenies from lineage tracing data using Cassiopeia. Genome Biology, 21 (92):1–27, 2020.

43. Geng Chen, John Paul Schell, Julio Aguila Benitez, Sophie Petropoulos, Marlene Yilmaz, Björn Reinius, Zhanna Alekseenko, Leming Shi, Eva Hedlund, Fredrik Lanner, Rickard Sandberg, and Qiaolin Deng. Single-cell analyses of X Chromosome inactivation dynamics and pluripotency during differentiation. Genome Research, 26(10):1342–1354, 2016.

44. Xiaojie Qiu, Andrew Hill, Jonathan Packer, Dejun Lin, Yi-An Ma, and Cole Trapnell. Single-cell mRNA quantification and differential analysis with Census. Nature Methods, 14(3): 309–315, 2017.

45. Georgi K Marinov, Brian A Williams, Ken McCue, Gary P Schroth, Jason Gertz, Richard M Myers, and Barbara J Wold. From single-cell to cell-pool transcriptomes: stochasticity in gene expression and RNA splicing. Genome Research, 24(3):496–510, 2014.

46. Dominic Grün, Lennart Kester, and Alexander van Oudenaarden. Validation of noise models for single-cell transcriptomics. Nature Methods, 11(6):637–640, 2014.

47. Christoph Ziegenhain, Beate Vieth, Swati Parekh, Björn Reinius, Amy Guillaumet-Adkins, Martha Smets, Heinrich Leonhardt, Holger Heyn, Ines Hellmann, and Wolfgang Enard. Comparative analysis of single-cell RNA sequencing methods. Molecular Cell, 65(4):631–643, 2017.

48. Huaiyu Mi, Anushya Muruganujan, and Paul D. Thomas. PANTHER in 2013: modeling the evolution of gene function, and other gene attributes, in the context of phylogenetic trees. Nucleic Acids Research, 41(D1):D377–D386, 2012.

49. David DeTomaso, Matthew G Jones, Meena Subramaniam, Tal Ashuach, Chun J Ye, and Nir Yosef. Functional interpretation of single cell similarity maps. Nature Communications, 10(4376), 2019.

50. Alexander Dobin, Carrie A Davis, Felix Schlesinger, Jorg Drenkow, Chris Zaleski, Sonali Jha, Philippe Batut, Mark Chaisson, and Thomas R Gingeras. STAR: ultrafast universal RNA-seq aligner. Bioinformatics, 29(1):15–21, 2013.

51. Kyle S Hubbard, Ian M Gut, Megan E Lyman, and Patrick M McNutt. Longitudinal RNA sequencing of the deep transcriptome during neurogenesis of cortical glutamatergic neurons from murine ESCs. F1000Research, 2(35):35, 2013.

52. Yu-Cheng T Yang, Chao Di, Boqin Hu, Meifeng Zhou, Yifang Liu, Nanxi Song, Yang Li, Jumpei Umetsu, and Zhi J Lu. CLIPdb: a CLIP-seq database for protein-RNA interactions. BMC Genomics, 16(51), 2015.

53. Philip J Uren, Emad Bahrami-Samani, Suzanne C Burns, Mei Qiao, Fedor V Karginov, Emily Hodges, Gregory J Hannon, Jeremy R Sanford, Luiz OF Penalva, and Andrew D Smith. Site identification in high-throughput RNA-protein interaction data. Bioinformatics, 28(23):3013–3020, 2012.

54. David Fischer, Fabian Theis, and Nir Yosef. Impulse model-based differential expression analysis of time course sequencing data. Nucleic Acids Research, 46:e119, 2018.

55. Yuhki Saito, Soledad Miranda-Rottmann, Matteo Ruggiu, Christopher Y Park, John J Fak, Ru Zhong, Jeremy S Duncan, Brian A Fabella, Harald J Junge, Zhe Chen, Roberto Araya, Bernd Fritzsch, A J Hudspeth, and Robert B Darnell. NOVA2-mediated RNA regulation is required for axonal pathfinding during development. eLife, 5, 2016.

56. Michaela Müller-McNicoll, Valentina Botti, Antonio M de Jesus Domingues, Holger Brandl, Oliver D Schwich, Michaela C Steiner, Tomaz Curk, Ina Poser, Kathi Zarnack, and Karla M Neugebauer. Sr proteins are nxf1 adaptors that link alternative rna processing to mrna export. Genes & Development, 30(5):553–566, 2016.

